# Disruption in A-to-I editing levels affects *C. elegans* development more than a complete lack of editing

**DOI:** 10.1101/273433

**Authors:** Nabeel S. Ganem, Noa Ben-Asher, Aidan C. Manning, Sarah N. Deffit, Michael C. Washburn, Emily C. Wheeler, Gene W. Yeo, Orna Ben-Naim Zgayer, Einav Mantsur, Heather A. Hundley, Ayelet T. Lamm

## Abstract

A-to-I RNA editing is widespread in eukaryotic transcriptomes and plays an essential role in the creation of proteomic and phenotypic diversity. Loss of ADARs, the proteins responsible for A-to-I editing, results in lethality in mammals. In *C. elegans*, knocking out both ADARs, ADR-1 and ADR-2, results in aberrant behavior and abnormal development. Studies have shown that ADR-2 can actively deaminate dsRNA while ADR-1 cannot. However, as most studies of *C. elegans* ADARs were performed on worms mutated in both ADAR genes, the effects observed cannot be attributed to a single ADAR or to the interactions between ADAR genes. Therefore, we set to study the effects of each *C. elegans* ADAR on RNA editing, gene expression, protein levels and the contribution of each of ADAR to the phenotypes observed in worms mutated in both genes, in order to elucidate their distinct functions. We found significant differences in the phenotypes observed in worms mutated in a single ADAR gene. Worms harboring *adr-1* mutations have a significant reduction in their lifespan, while worms harboring *adr-2* mutations have extended lifespan. We also observed severe abnormalities in vulva formation in *adr-1* mutants, and we suggest that these phenotypes are a result of an RNA editing independent function of ADR-1. Mutations in each ADAR resulted in expressional changes in hundreds of genes, and a significant downregulation of edited genes. However, very few changes in the protein levels were observed. In addition, we found that ADR-1 binds many edited genes and primarily promotes editing at the L4 stage of development. While editing still occurs in the absence of ADR-1, most of the editing occurs in genes that are edited in wildtype worms, suggesting that ADR-1 does not prevent editing by binding to and protecting the RNA but rather enhances or promotes editing. Our results suggest that ADR-1 plays a significant role in the RNA editing process and by altering editing levels it causes the severe phenotypes that we observed. In contrast, a complete lack of RNA editing is less harmful to the worms. Furthermore, our results indicate that the effect of RNA editing on the protein content in the cell is minor and probably the main purpose of these modifications is to antagonize or enhance other gene regulatory mechanisms that act on RNA.

## Introduction

A-to-I RNA editing is a conserved process in which Adenosine deaminases that act on RNA (ADAR) enzymes convert adenosine to inosine within double-stranded regions of RNA (Bass, 2006). Inosine is read as guanosine by the translation machinery and thus can change protein amino acid content and function. Several examples of protein changes in the human brain were described (Burns et al., 1997, Pullirsch and Jantsch, 2010, Werry et al., 2008), however the majority of editing sites in humans are in non-coding regions of the transcriptome (mainly in *Alu* repeats) (Athanasiadis et al., 2004, Barak et al., 2009, Blow et al., 2004, Kim et al., 2004, Levanon et al., 2004, Li et al., 2009b). In mammals, ADARs are essential – knockout mice are either embryonic lethal (ADAR1) or lethal shortly after birth (ADAR2) (Wang et al., 2000, Higuchi et al., 2000). However, in model organisms such as *C. elegans* and *D. melanogaster*, strains that lack all RNA editing are viable, although they exhibit behavioral and anatomical defects (Tonkin et al., 2002, Palladino et al., 2000).

Many A-to-I editing sites have been identified in the *C. elegans* transcriptome using high-throughput RNA sequencing, and most of them are located in introns and other non-coding regions (Goldstein et al., 2017, Whipple et al., 2015, Wu et al., 2011, Zhao et al., 2015). RNA editing in *C. elegans* is developmentally regulated. The overall frequency of RNA editing is higher in embryos than in other stages (Zhao et al., 2015). However, there are genes that undergo frequent RNA editing at the L4 developmental stage and almost no editing at the embryo stage, despite the transcripts being expressed to a similar level in both stages (Goldstein et al., 2017). This points to the fact that the editing activity in *C. elegans* is highly regulated.

*C. elegans* possess two ADAR genes: *adr-1* and *adr-2*. Both proteins share the common ADAR enzyme structure: highly conserved C-terminal deaminase domain and variable number of N-terminal dsRNA binding motifs (two dsRBD in ADR-1, and one in ADR-2, Supp Figure 1) (Tonkin et al., 2002). However, ADR-2 is the only active adenosine deaminase in *C. elegans*, as knockout of *adr-2* abolishes all A-to-I RNA editing (Washburn et al., 2014). On the other hand, ADR-1 acts as a regulator of ADR-2, regulating editing efficiency by interacting with ADR-2 and with ADR-2 targets through its dsRNA binding domains (Rajendren et al., 2018, Washburn et al., 2014). Expression analysis of GFP-tagged ADR-1 revealed that it is expressed mainly in the nervous system and developing vulva, but expression occurs in all developmental stages (Tonkin et al., 2002). Worms harboring deletions in both *adr-1* and *adr-2* genes are viable but have been reported to exhibit phenotypes including chemotaxis defects, altered lifespan, and decreased expression of transgenes (Knight and Bass, 2002, Sebastiani et al., 2009, Tonkin et al., 2002). Worms harboring a specific mutated *adr-1* allele were also shown to have a mild protruding-vulva (Pvl) phenotype (Tonkin et al., 2002).

Although thousands of editing sites were found in *C. elegans* (Goldstein et al., 2017, Whipple et al., 2015, Wu et al., 2011, Zhao et al., 2015), most of these sites are located in non-coding regions and therefore are hard to link to the phenotypes of ADAR mutant worms. A slight general reduction in expression of *C. elegans* genes that undergo editing in 3’ untranslated regions (UTRs) occurs in worms harboring deletions in both *adr-1* and *adr-2* (Goldstein et al., 2017); however, most of these changes were not directly linked to phenotypic consequences. Recently, it was shown that a gene, *clec-41*, which is edited in the 3’ UTR is downregulated in neural cells lacking *adr-1* or *adr-2*. Furthermore, the reduction in *clec-41* contributes to the chemotaxis defects of worms lacking *adr-2* (Deffit et al., 2017). Interestingly, mutations in RNAi genes in worms harboring mutations in *adr-1* and *adr-2* can rescue the chemotaxis and lifespan phenotype of the worms (Sebastiani et al., 2009, Tonkin and Bass, 2003), implicating RNAi involvement in the function of RNA editing. However, the specific transcripts that are important for these phenotypes and are altered by RNAi are unknown. Furthermore, it is still unknown what the main function of RNA editing is in *C. elegans*. There were some predictions of changes in protein structure (Goldstein et al., 2017, Zhao et al., 2015) caused by editing in coding regions of genes, though this was shown on a handful of genes and the changes in protein structure were not validated. A new function for RNA editing emerged from studies in mammals, in which mammalian ADAR1 was implicated in preventing aberrant activation of the innate immune response from self produced dsRNAs (George et al., 2016, Liddicoat et al., 2015, Mannion et al., 2014). It is possible that in *C. elegans* RNA editing has a similar function to prevent protection mechanisms such as RNAi to attack self-produced RNA (Ganem and Lamm, 2017, Reich et al., 2018). In this case the phenotypes observed in ADAR mutants might result from changes in the RNA level and not necessarily in the protein level.

Many of the studies done so far on *C. elegans* examined phenotypes, editing levels, and expression changes in worms harboring mutations in both *adr-1* and *adr-2* genes (for example (Goldstein et al., 2017, Tonkin et al., 2002, Whipple et al., 2015, Wu et al., 2011, Zhao et al., 2015)). While some of these previous studies indicated that lack of either ADAR genes resulted in chemotaxis and lifespan defects (Sebastiani et al., 2009, Tonkin et al., 2002), ADR-1 probably affects the expression and function of edited genes and the severity of the phenotypes observed in a different manner than ADR-2. In addition, ADAR genes might have functions beyond RNA editing. For example, in mice ADARs were shown to affect splicing independent of their editing functionality (Solomon et al., 2013) and ADAR1 was found to inhibit Staufen mediated mRNA decay, independently of its A-to-I editing function (Sakurai et al., 2017). To understand the contribution of each of the *C. elegans* ADAR genes to the RNA editing process, gene regulation, and the phenotypes exhibited in ADAR mutant worms, we performed a comprehensive phenotypic, transcriptomics and proteomics analysis on worms harboring mutations in *adr-1* or *adr-2* separately. We found that the phenotypes observed in *adr-1* mutants are more severe and some are distinct from those we observed in *adr-2* mutants. We found that both genes affect the expression of edited genes but the effect of ADR-1 is much more prominent. Editing still occurs in *adr-1* mutants; however, the number of editing sites is reduced significantly at the L4 developmental stage. In addition, edited genes are also a significant portion of genes bound by ADR-1 either directly or indirectly. These results implicate ADR-1 as an important component of the RNA editing process and that changes in the levels of editing cause more developmental defects than a complete lack of editing. Our results also suggest that the main function of ADARs and RNA editing is to regulate RNA expression and not the protein content in a cell.

## Results

Previous studies suggested that ADR-1 is not necessary for the editing process and is mostly a regulator of ADR-2 activity (Rajendren et al., 2018, Tonkin et al., 2002, Washburn et al., 2014). Editing is still observed in *adr-1* mutants although at different levels (Tonkin et al., 2002, Washburn et al., 2014). To explore the function of *adr-1* and *adr-2* in the editing process separately, we performed phenotypic, transcriptomics and proteomics analyses on two deletion mutations for each gene. Both deletions of the *adr-2* gene are expected to result in a non-functional protein, removing either the deaminase domain or most of the protein (Supp Figure 1, Supp Figure 2, (Tonkin et al., 2002)). As expected, we did not observe editing in mutants containing either one of these deletions (Supp Figure 3,4). ADR-1 has five annotated isoforms (Supp Figure 1). Two isoforms are predicted to give rise to truncated proteins that do not contain either the dsRNA binding domains or the deaminase domain. The dsRNA binding domains in ADR-1 were shown to be required to regulate editing by ADR-2 (Rajendren et al., 2018, Washburn et al., 2014) and both available deletions completely delete or disrupt the dsRNA binding domains (Tonkin et al., 2002, Washburn et al., 2014). Indeed, we observed changes in the editing levels in both deletion mutants and as expected editing was not completely abolished (Supp Figure 3,4).

### *adr-1* mutant phenotypes are more severe than those observed in *adr-2* mutants

Worms harboring mutations in either or both *adr-1* and *adr-2* were previously shown to have reduced life span and chemotaxis defects (Sebastiani et al., 2009, Tonkin et al., 2002). We therefore sought to explore if both genes contribute equally to these phenotypes. If these phenotypes are a result of abolished editing, we expected to observe the same phenotypes in *adr-2* mutants and to a lesser extent in *adr-1* mutants. Chemotaxis experiments that were done by Tonkin et al (Tonkin et al., 2002) were repeated, but with two deletion mutations for each of the genes instead of one. We received similar results that all mutants have chemotaxis defects (Supp Figure 5). The chemotaxis defects were restored to normal in transgenic worms harboring FLAG-ADR-1 in an *adr-1* mutant background (Washburn et al., 2014) (Supp Figure 5). These results are in line with a study that found that both ADR-1 and ADR-2 affect expression of the *clec-41* gene in neural cells and overexpressing *clec-41* in *adr-2* mutants rescues the chemotaxis defect (Deffit et al., 2017). Interestingly, we observed an upregulation of *clec-41* in *adr-1* mutant worms in embryo stage but not in L4 stage and not in *adr-2* mutants (see below). This upregulation was also observed previously in L1 worms (Deffit et al., 2017). However, in neural cells *clec-41* was shown to be downregulated in *adr-1* mutants (Deffit et al., 2017), suggesting that ADR-1 and ADR-2 may provide tissue-specific gene regulation in addition to development-specific gene regulation.

In addition, we performed life span experiments on worms with deletions in both *adr* genes and worms with a deletion in a single *adr* gene. Strains harboring mutations in both *adr-1* and *adr-2* had reduced life span or similar life span to wildtype worms (Figure 1A). Surprisingly, we found that both deletion strains of *adr-1* significantly reduce life span, while both strains with deletions in *adr-2* significantly extend the life span compared to wildtype worms (Figure 1A). Strains carrying the extra chromosomal array FLAG-ADR-1 in an *adr-1* deletion background were able to slightly rescue the life span phenotype of *adr-1* mutation (Figure 1B). Interestingly, strains carrying a FLAG-ADR-1 with mutation in the dsRNA binding domains (dsRBM) were able to rescue the life span phenotype as well as wild-type ADR-1, suggesting that either the ability of ADR-1 to promote life span is independent of dsRNA binding or possibly mutant ADR-1 still has the ability to bind mRNAs other than the edited mRNAs that were previously shown to have disrupted ability to interact with the ADR-1 dsRBM mutant (Washburn et al., 2014).

**Figure 1.**
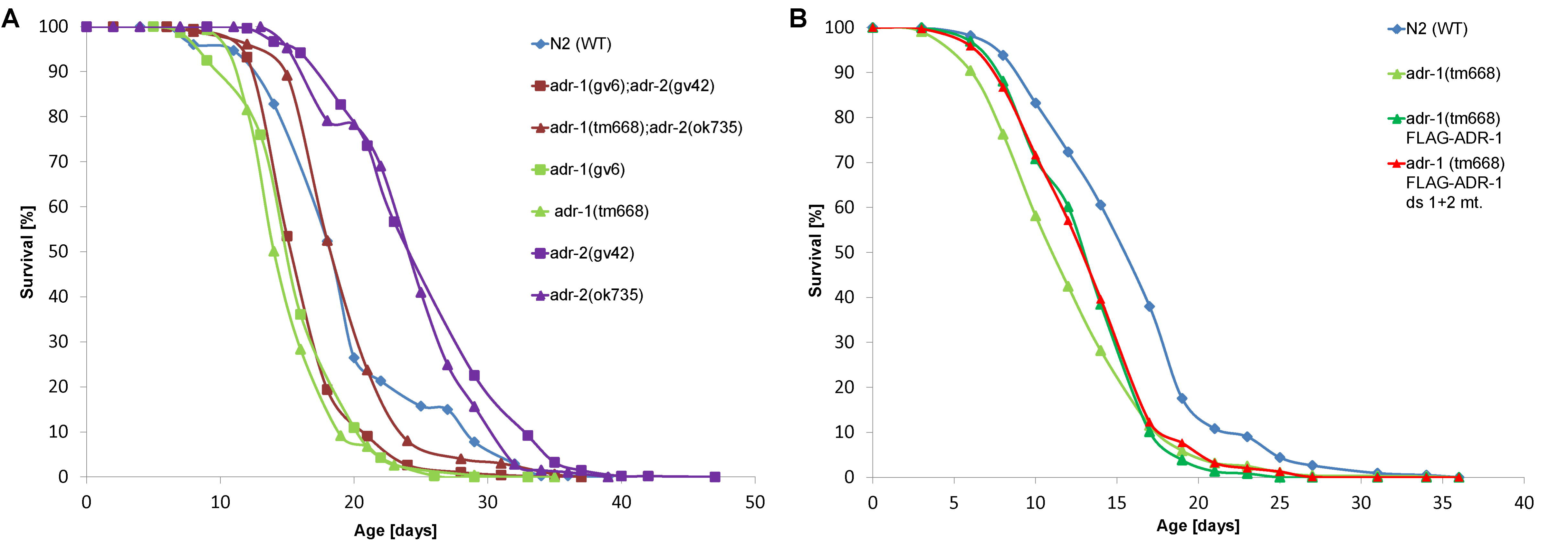
*adr-1* and *adr-2* mutants have opposite lifespan phenotypes. (A) The lifespan of the mutant worms was followed and the mean survival curves are presented for N2 (WT), *adr-1*(gv6)I, *adr-1*(tm668)I, *adr-2*(gv42)III, *adr-2*(ok735)III, *adr-1*(gv6)I; *adr-2*(gv42)III and *adr-1*(tm668)I; *adr-2*(ok735)III. Mutations in *adr-1* gene (*adr-1*(gv6)I or *adr-1*(tm668)I) reduce the lifespan of the worm comparing to the WT worms, while mutation in *adr-2* gene (*adr-2*(gv42)III or *adr-2*(ok735)III) extend the lifespan of the worm comparing to the WT worms. Mutants for both genes have the same pattern as the WT worms. (B) The mean survival curves are presented for WT, *adr-1* (tm668), and the rescue strains *adr-1* (tm668) FLAG-ADR-1: *adr-1*(tm668) I blmEx1[3XFLAG-*adr-1* genomic, *rab3*::*gfp*::*unc-54*] and *adr-1* (tm668) FLAG-ADR-1 ds1+2 mutant: *adr-1*(tm668) I blmEx1[3XFLAG-*adr-1* genomic with mutations in dsRBD1 (K223E, K224A, and K227A) and dsRBD2 (K583E, K584A and K587A), *rab3*::*gfp*::*unc-54*]. Both transgenic *adr-1* rescue strains have extended lifespan than *adr-1*(tm668). Each experiment was repeated at least five times.

Previously *adr-1* was shown to be highly expressed in the vulva and one of the *adr-1* deletion strains exhibited a slight protruding-vulva (Pvl) phenotype (Tonkin et al., 2002). To study if this phenotype is strain specific because of a second mutation in a close gene or because of lack of *adr-1*, we counted the fraction of worms with Pvl phenotype in all strains. We found a significant Pvl phenotype in all mutant strains compared to wildtype strains although the phenotype fraction from total worms was very low (less than 10% in all strains) (Figure 2A). ADR-1 mutants had the highest Pvl phenotype fraction, which was significantly reduced in FLAG-ADR-1 and FLAG-ADR-1 with dsRBM mutations transgene worms (Figure 2A). We conclude that all ADAR mutants have developmental phenotypes although ADR-1 seems to be more important for normal development than ADR-2.

**Figure 2.**
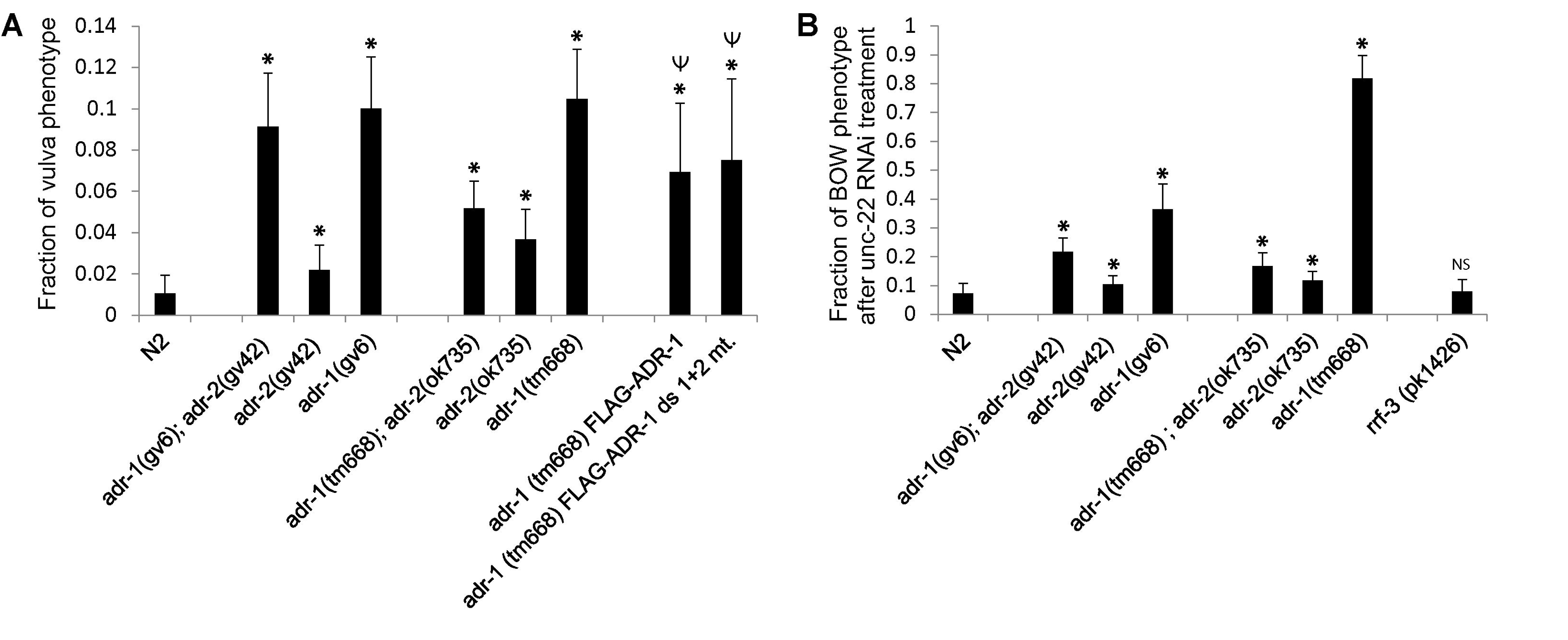
*adr-1* mutants have high frequency of vulva abnormalities. (A) ADAR mutant strains were scored for pvl phenotype and the fraction of worms presenting the phenotype from total worms is presented. P-value was calculated by two samples unequal variance heteroscedastic *t*-test; * comparing to WT; *P*<0.01; Ψ comparing to *adr-1*(tm668)I); *P*<0.01. (B) Worms were subjected to *unc-22* RNAi and the fraction of worms presenting Bag of worms phenotype is presented. P-value was calculated by two samples unequal variance heteroscedastic *t*-test; *comparing to WT; *P*<0.01. Each experiment was repeated at least three times and the standard deviation is shown by error bars.

We previously suggested that downregulation of genes in ADAR mutants could be a consequence of the antagonistic relationship between RNA editing and RNAi (Goldstein et al., 2017). The possibility that ADAR genes and RNA editing itself are antagonistic to the RNAi process was raised previously as editing and RNAi both involve dsRNA substrates, transgene silencing was observed in ADAR double mutants, and changes in the amount of siRNAs generated in wild-type worms and those lacking both ADARs (Goldstein et al., 2017, Knight and Bass, 2002, Reich et al., 2018, Tonkin and Bass, 2003, Warf et al., 2012, Whipple et al., 2015, Wu et al., 2011). Hypersensitivity to exogenous RNAi is a phenotype related to the ERI/RRF-3 endogenous RNAi pathway (Simmer et al., 2002), which is also reflected by transgene silencing. Therefore, hypersensitivity to RNAi was tested before in several ADAR mutants (Knight and Bass, 2002, Ohta et al., 2008), which did not show hypersensitivity. We tested hypersensitivity to RNAi in all ADAR single and double mutants by triggering exogenous RNAi by feeding the worms with bacteria producing dsRNA against *lin-1* or *unc-22* genes (Simmer et al., 2002) and scoring the phenotype. We did not observe high enrichment of the multivulva phenotype in all ADAR mutant worms as compared to wildtype when triggering RNAi against *lin-1* (Supp Figure 6). As expected, *rrf-3* mutant worms, which are hypersensitive to RNAi have a significant high fraction of the phenotype (Supp Figure 6). Triggering *unc-22* RNAi we expected a twitching phenotype, while observing a strong twitching phenotype we also noticed a new phenotype of bag of worms (Figure 2B). The bag of worms phenotype was observed in a high fraction in *adr-1* mutant worms to a lesser extent in the *adr-1*;*adr-2* double mutant and in *adr-2* mutants. This phenotype is not a result of hypersensitivity to RNAi because it was not enriched in *rrf-3* mutants (Figure 2B). We conclude that this bag of worm phenotype and the pvl phenotype are specific to *adr-1* and might be a result of a different function of ADR-1 distinct from RNA editing.

### Both ADR-1 and ADR-2 affect expression of genes edited at their 3’UTR

Previously we demonstrated that the expression of genes with edited 3’UTRs is slightly reduced in worms harboring deletions in both ADAR genes (Goldstein et al., 2017). The list of genes that are edited at their 3’UTR was based on a screen for editing sites in non-repetitive regions in the transcriptome. In total 77 genes with 3’UTR edited sites were identified (Goldstein et al., 2017). However, many edited sites were not included in the list even though they are in a very close proximity to the 3’UTR annotation of genes. These include genes that were biochemically identified by others to be edited at their 3’UTR for example, *alh-7* and *C35E7.6* (Morse et al., 2002, Morse and Bass, 1999). To extend the list of genes edited at their 3’UTR, we manually annotated the edited sites in non-repetitive regions that are in proximity to genes. We added to the list genes where multiple editing sites are in the same orientation as the gene and in very close proximity to their annotated 3’UTR or within the 3’UTR based on the newest version of Wormbase (see material and methods). Overall, we added 59 genes and the list now includes 136 genes (Supp Table 1). To re-examine the conclusion regarding expression levels of genes edited in 3’UTRs in worms harboring deletions in both *adr-1* and *adr-2*, we reanalyzed the expression data from (Goldstein et al., 2017). We observed similar results with the 136 genes, their expression levels were slightly reduced compared to all genes in both embryo and L4 developmental stages (p-value < 0.003 calculated by Welch two-sample T-test analysis for genes with Padj value < 0.05), suggesting that the newly identified genes edited in 3’UTRs are *bona fide* ADAR regulated genes (Supp Figure 7).

To study if the expression levels of 3’UTR edited genes are affected in the single deletion mutants, we generated RNA-seq data from 3 biological replicates of each of the strains and the wildtype strain in embryo and L4 developmental stages. We observed the tendency for reduced expression of 3’UTR edited genes in all *adr-1* and *adr-2* mutants in embryo stage (Figure 3A, B, p-value < 2.2e-16 for *adr-1 (gv6)*, p-value = 0.002 for *adr-1 (tm668)*, p-value < 0.002 for *adr-2 (gv42)*, and p-value = 2.5e-05 for *adr-2 (ok735)*). However, the expression of 3’UTR edited genes does not change in all single ADAR mutants in L4 developmental stage (Figure 3C, D). This is in contrast to what we observed before in the double mutants (Goldstein et al., 2017). Thus, the effect of ADAR mutations on the expression of 3’UTR edited genes is stronger in embryo than in L4 developmental stage.

**Figure 3.**
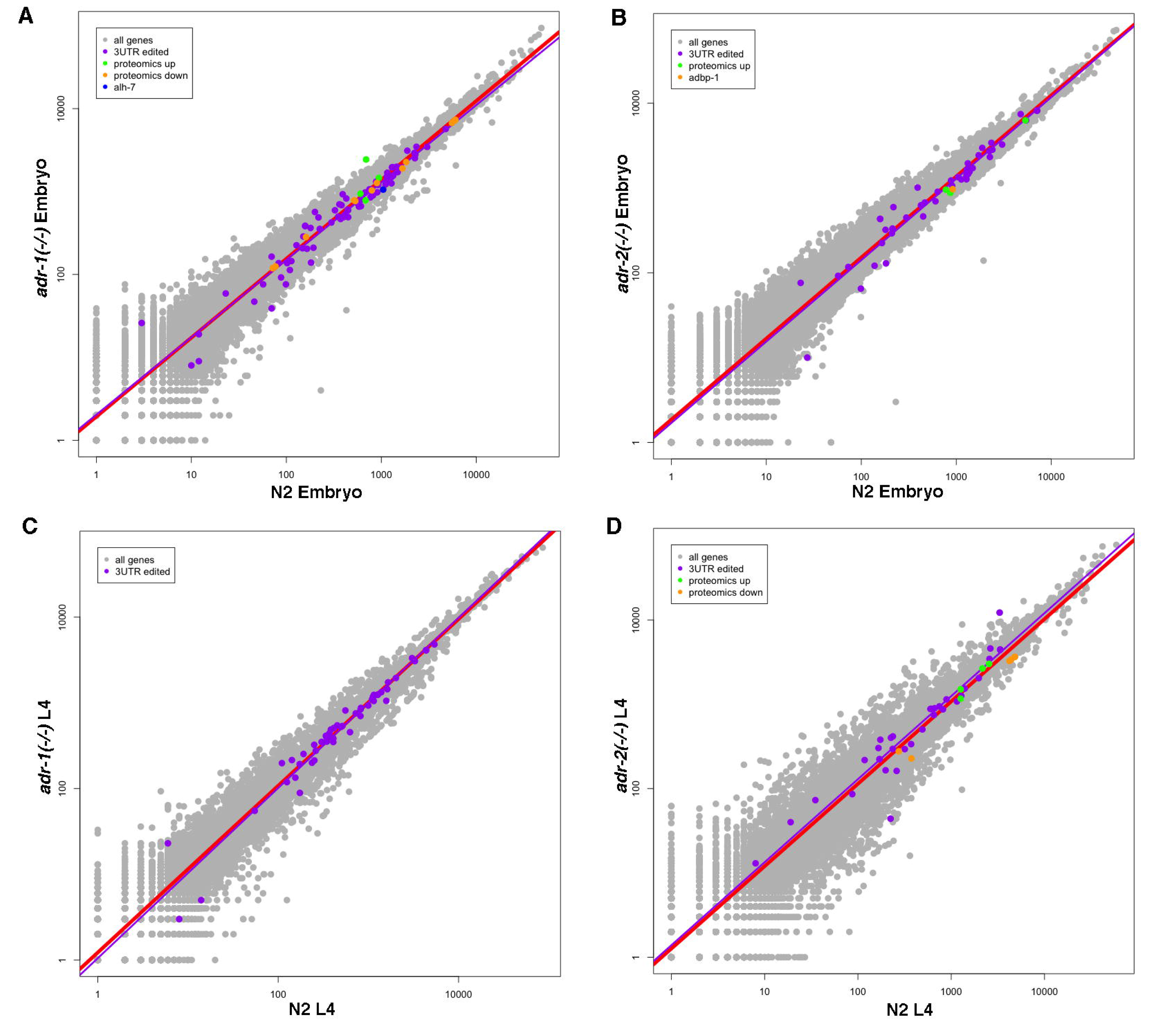
Genes edited at their 3’UTR are downregulated in *adr-1* and *adr-2* mutants at the embryo stage. Log scale plots presenting gene expression in wildtype (N2) worms versus *adr-1* mutant worms (A, C) or *adr-2* mutant worms (B, D) at embryo stage (A, B) and L4 stage (C, D). Every dot in the graphs represents a gene. Red line is the regression line for all genes. 3’UTR edited genes with significant padj value are in purple and their regression line is presented in purple. Downregulated genes found by the proteomics analysis are in orange and upregulated genes are in green. *Alh-7* gene is downregulated in *adr-1* mutants at embryo stage by all analyses (blue in A). *Adbp-1* gene is downregulated in *adr-2* mutants by all analyses at embryo stage (orange in B).

### ADR-1 affects the expression of edited genes

To further explore the expression of genes in *adr-1* and *adr-2* mutants, we identified the genes with a 2-fold or greater change in expression and a significant adjusted p-value after Benjamini-Hochberg correction in each of the single ADAR mutants in each developmental stage. We further shortened the list by including only genes that were significantly differentially expressed in both deletion mutations for each gene (Supp Tables 2,3,4) to avoid allele specific background effects on gene expression. We found more upregulated than downregulated genes in both ADAR mutants in all developmental stages (Supp Table 4) with significant overlap (p-value < 0.01 calculated by hypergeometric distribution and by chi-square test) of differential expressed genes between embryo and L4 developmental stages (Supp Figure 8). There is also a very significant overlap (p-value < e^−5^) between differentially expressed genes in *adr-1* mutants and *adr-2* mutants in all stages (Supp Figure 9). However, 3’UTR edited genes were not a substantial part of the differentially expressed genes in either ADAR mutants (p-value not significant, Supp Figure 9). To explore if genes edited in regions other than the 3’UTR are differentially expressed in ADAR mutants, we compared the genes differentially expressed in each ADAR mutant and genes edited at the embryo or L4 developmental stage (described in (Goldstein et al., 2017)). We did not find a significant overlap between differentially expressed genes in *adr-2* mutants and genes edited at L4 or at embryo stage (non-significant p-value, Supp Figure 10). Surprisingly, a significant portion of the upregulated genes in *adr-1* mutants are genes edited at the L4 stage (p-value < 0.01, Figure 4). Downregulated genes in *adr-1* mutants are not enriched for edited genes (non-significant P-value, Figure 4). These results suggest that *adr-1* not only affects the level of editing but also the expression of edited genes.

**Figure 4.**
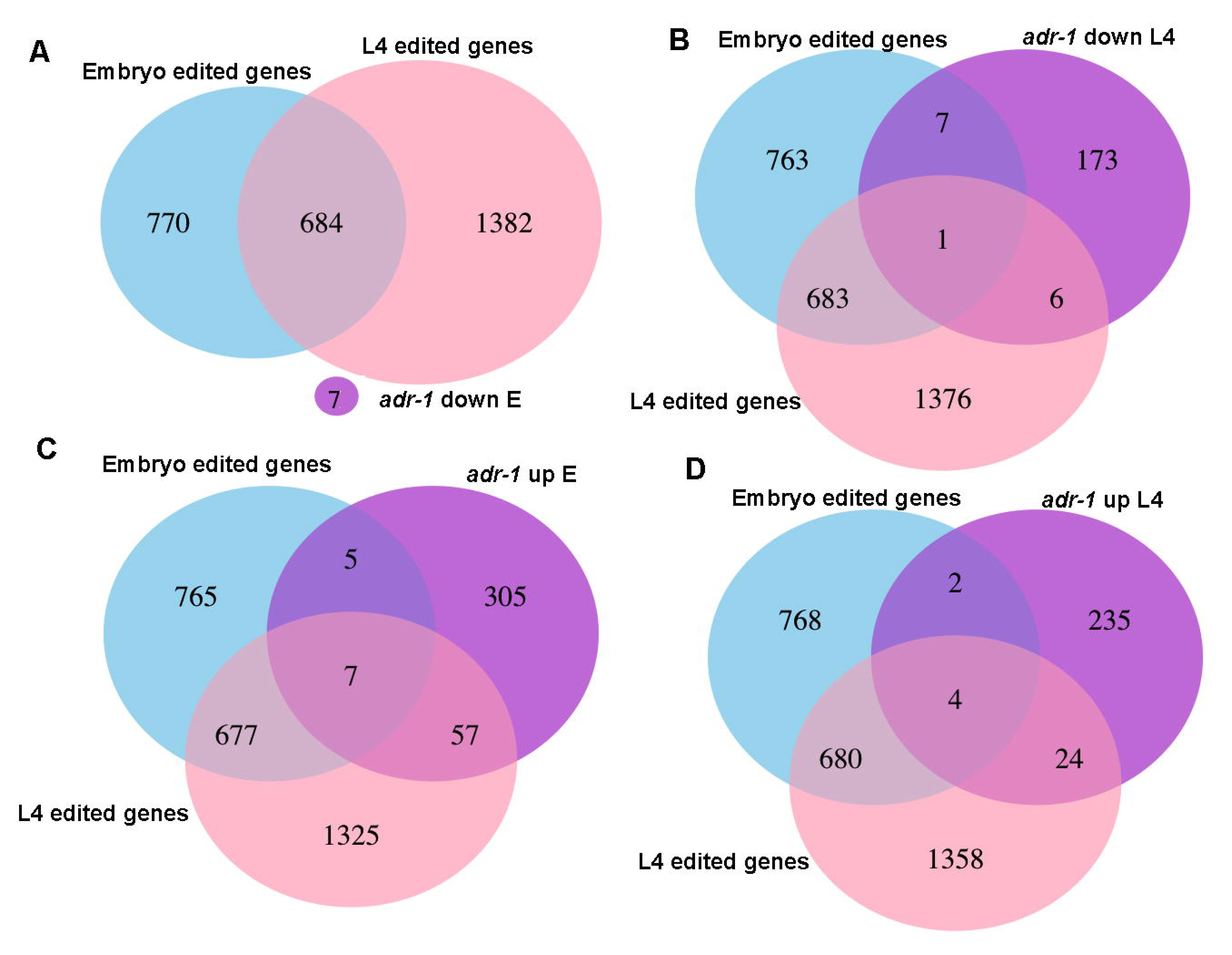
Significant portions of genes that are edited at L4 stage are upregulated in *adr-1* mutants. Venn diagrams presenting the intersections between edited genes at embryo or L4 developmental stages and A. Genes that their expression is downregulated at embryo stage in *adr-1* mutants. B. Genes that their expression is downregulated at L4 stage in *adr-1* mutants. C. Genes that their expression is upregulated at embryo stage in *adr-1* mutants. D. Genes that their expression is upregulated at L4 stage in *adr-1* mutants.

### ADR-1 role in regulating RNA editing is stronger at the L4 stage

The increased expression of edited genes and the alterations in editing levels in *adr-1* mutants suggests that ADR-2 editing of these genes is assisted by ADR-1 and may be important to stabilize gene expression. As for the lack of connection between the downregulated genes in the *adr-1* mutant worms and editing in wildtype worms, it is possible that these downregulated genes contain editing sites in the *adr-1* mutants and not in wild-type worms. This would suggest that ADR-1 also has a role in binding RNA and protecting or preventing the RNA from editing by ADR-2. Therefore, new edited sites, “the protected sites”, should emerge in the high-throughput sequencing datasets obtained in the absence of ADR-1. To test this hypothesis, we performed a screen to identify editing sites in *adr-1* mutants similarly to (Goldstein et al., 2017), with the difference that only nucleotide changes that appeared in both *adr-1* mutant alleles were considered. When counting the number of nucleotide changes that were identified in *adr-1* mutants, surprisingly we found a significant reduction in the amount of editing sites identified in *adr-1* mutants at L4 developmental stage compared to embryo (Figure 5A, P-value < 0.01 calculated by Fisher Exact test, Supp Table 5). Next, we tested whether there are editing sites in *adr-1* mutants that are not present in wildtype worms by comparing the lists of nucleotide changes. While the number of editing sites that are only present in *adr-1* mutants was not above the background of the other nucleotide changes at the L4 stage, we found an enrichment of editing sites that are present only in the *adr-1* mutants at the embryo stage (Figure 5A, Supp Table 5). To further study these sites, we compared genes with editing sites that are only edited in *adr-1* mutants at the embryo stage to edited genes found in wildtype worms. We found that most of the genes with editing sites unique to *adr-1* mutants have other sites that are edited in wildtype worms (Figure 5B). In addition, these genes do not have a significant change in expression in *adr-1* mutants (Figure 5 C, D). Overall, these results suggest that ADR-1 does not protect the dsRNA from being edited by ADR-2 but rather directs and enhances ADR-2 editing especially at the L4 developmental stage. This is in line with our findings that many genes that are upregulated in *adr-1* mutants are only edited at the L4 developmental stage (Figure 4).

**Figure 5.**
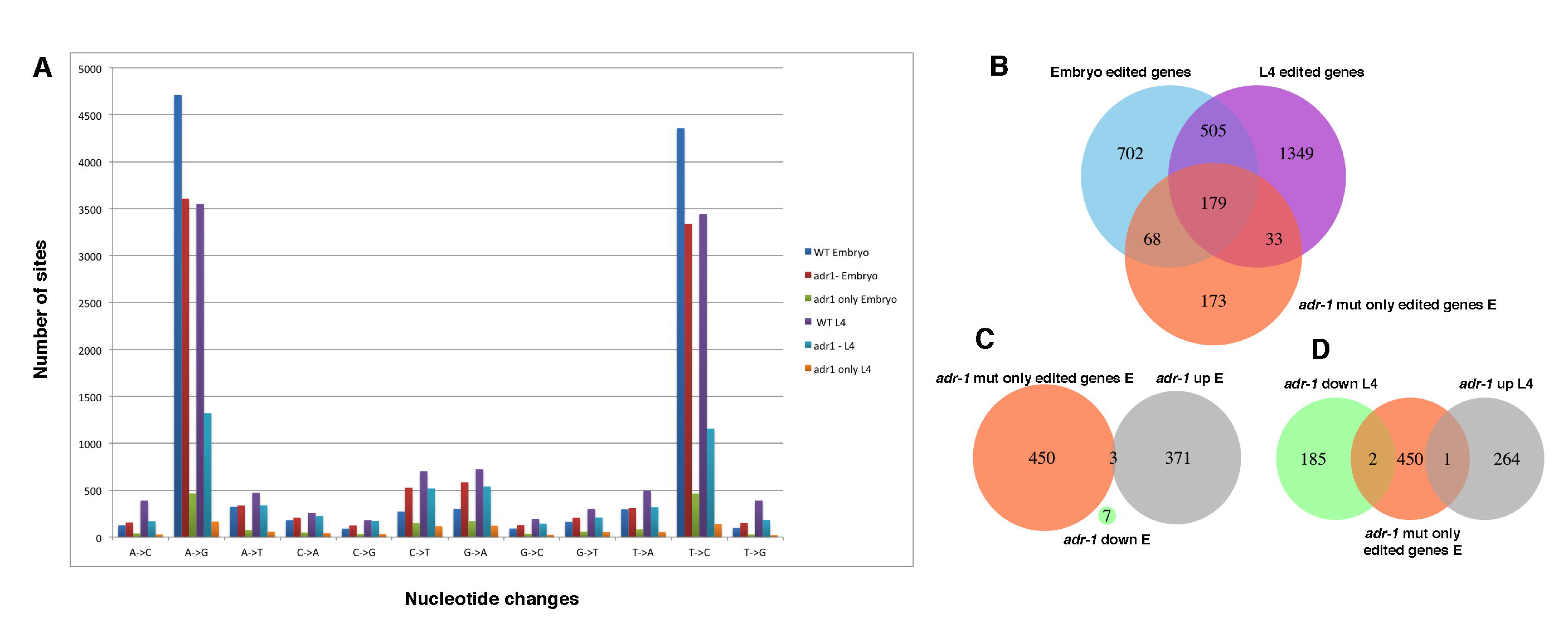
Most of the editing sites that appear only in *adr-1* mutants are in genes with editing sites in wildtype worms. (A) A bar graph representing nucleotide changes found in wildtype and *adr-1* mutant worms at L4 and embryo developmental stages. Also presented are nucleotide changes that were found in *adr-1* mutants but not in wildtype worms. (B-D) Venn diagrams presenting the intersection between genes with editing sites that were detected in *adr-1* mutants and not in wildtype and (B) Genes edited in wildtype worms at embryo or L4 developmental stages. (C). Genes that their expression is upregulated or downregulated at embryo stage in *adr-1* mutants. (D) Genes that their expression is upregulated or downregulated at L4 stage in *adr-1* mutants.

### ADR-1 binds edited genes

To study if ADR-1 affects the expression of genes by binding their RNA, we performed RNA immunoprecipitation assay on young adult worms using the FLAG antibody and worms expressing the FLAG-1:ADR-1 transgene or worms lacking *adr-1* as a negative control. The bound RNAs were extracted, polyA selected and sequenced using high-throughput sequencing (RIP-seq, Supp Table 6). Almost one-third of the 3’UTR edited genes are bound by ADR-1 (Supp Figure 11) and a significant portion of edited genes in general as determined by hypergeometric distribution test (p-value < e^−44^, Figure 6). These results confirm that ADR-1 regulates edited genes by binding their RNA either directly or through a common interacting RNA binding protein. Interestingly, we found that ADR-1 binds *unc-22*, which has an important role in the regulation of the actomyosin dynamics (Benian et al., 1989), and *unc-54*, which encodes a muscle myosin class II heavy chain (Waterston, 1989). Mutations in *unc-54* can suppress the twitching phenotype of *unc-22*, which suggests an interaction between these genes (Moerman et al., 1982). Thus, the new bag of worms phenotype that we observed when *adr-1* mutant worms were subjected to *unc-22* RNAi (Figure 2B) suggests a function for ADR-1 in muscle formation. This function is probably distinct from the function of ADR-1 in RNA editing as we did not find editing in *unc-54* RNA and *unc-22* only has one editing site in an intron. In addition, this phenotype is not as significant in *adr-2* mutants.

**Figure 6.**
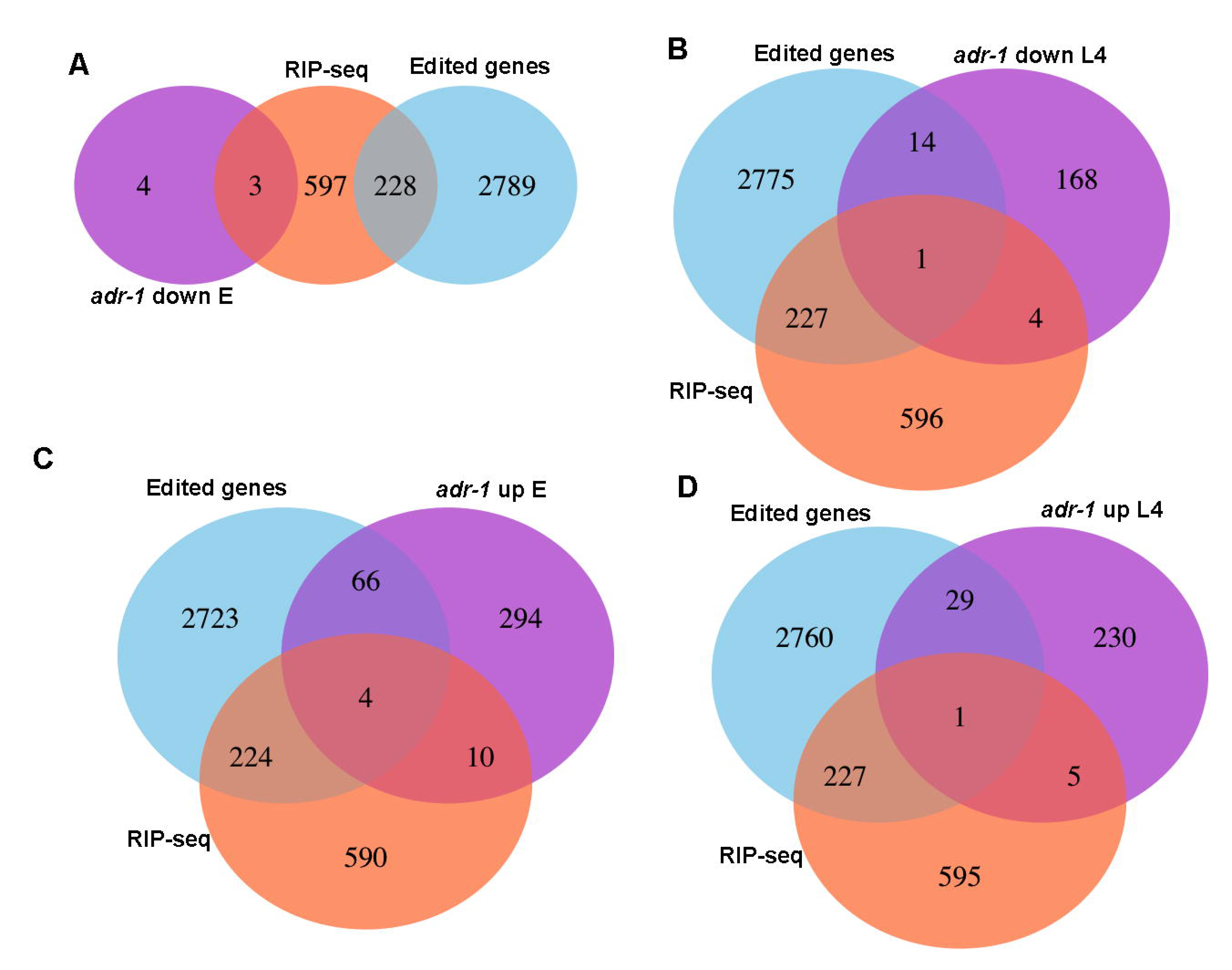
ADR-1 binds edited genes. Venn diagrams presenting the intersections between edited genes, genes identified as bound by ADR-1 using RIP-seq analysis and A. Genes that their expression is downregulated at embryo stage in *adr-1* mutants. B. Genes that their expression is downregulated at L4 stage in *adr-1* mutants. C. Genes that are upregulated at the embryo stage in *adr-1* mutants. D. Genes that are upregulated at the L4 stage in *adr-1* mutants.

### RNA editing does not affect the protein levels of edited genes

To study if the changes in RNA levels of genes in ADAR mutants affect their protein levels as well, we performed a comparison between the proteome content of wildtype worms and *adr-1* or *adr-2* mutant worms at embryo (Supp Table 7) or L4 developmental stage (Supp Table 8). We extracted proteins from three biological replicates of each strain, trypsinized the proteins and quantified them using mass spectrometry (see material and methods). These experiments identified 5984 proteins in total, 1426 were only identified in the embryonic samples and 1333 were specific to the L4 stage. While there was a significant representation of edited genes in the proteomics analysis (Supp Figure 12, P-value < e-4 for all groups of edited genes), the protein levels of only 22 genes were significantly changed in ADAR mutants (Supp table 9). Out of the 22 genes where protein levels changed in both *adr-1* mutant strains or in both *adr-2* mutants strains (not including ADAR genes themselves), 4 genes were also found to be edited, *C06A5.6*, *W07G4.3*, *Y54E2A.4*, and *alh-7*. These results are not surprising as most of the editing sites are in non-coding regions (Goldstein et al., 2017) and many of the edited genes in human and *C. elegans* as well are probably not protein coding genes. Interestingly, *alh-7*, which undergoes editing at its 3’UTR and is a highly conserved neuronal gene (Morse et al., 2002), was significantly downregulated in *adr-1* mutants both at the RNA level and the protein level at embryo stage (Figure 3). Another interesting gene is *adbp-1*, which is a regulator of *adr-2* (Ohta et al., 2008). ADBP-1 was shown to interact with ADR-2 and facilitates its cellular localization (Ohta et al., 2008). We found that the protein level of *adbp-1* was significantly downregulated in both *adr-2* mutants but not the mRNA levels (Figure 3, Supp Tables 2,6), while ADR-2 protein was also significantly downregulated in *adbp-1* mutant (Supp Figure 13). These results suggest that the regulation between ADR-2 and ADBP-1 is not unidirectional but both proteins regulate each other’s stability.

## Discussion

In this work we performed a comprehensive phenotypic, transcriptomic and proteomic analysis on the two ADAR genes in *C. elegans* to explore their individual role in the RNA editing process. We confirmed that ADR-2 is the only active enzyme, but found that ADR-1 regulates not only the editing process, but also the expression of edited genes. This comprehensive analysis was performed on two different deletion alleles for each ADAR gene to rigorously identify effects of each ADAR and avoid allele specific bias.

### ADR-1 has distinct phenotypes, and likely distinct functions from ADR-2

The analysis revealed several interesting abnormal phenotypes, some of which are specific to *adr-1* mutants. The aberrant chemotaxis phenotype that was previously described (Tonkin et al., 2002) was apparent in all ADAR mutants (supp Figure 5) and was rescued by an ADR-1 transgene. The expression of the edited gene *clec-41* in neuronal cells was shown to be important for this phenotype (Deffit et al., 2017). This suggests that the absence or changes in the levels of editing in specific genes causes the aberrant chemotaxis phenotype. The decrease in the lifespan was previously observed in worms harboring mutations in either *adr*-1 or *adr-2* genes or both (Sebastiani et al., 2009). When we examined lifespan of worms harboring the single mutants, we found that the two ADAR genes contribute to the lifespan of the worms in the opposite direction. The changes in editing levels in ADAR mutants might produce these phenotypes. One possibility is when editing is absent in *adr-2* mutants the worms live longer and when editing levels are reduced or even elevated in specific genes in *adr-1* mutants it causes a reduction in lifespan. As a decrease in the lifespan is a very common phenotype in *C. elegans* and reduced expression of many edited genes may cause a similar phenotype (Goldstein et al., 2017, Zhao et al., 2015), it is hard to pinpoint a particular gene that underlies this phenotype. We only observed a partial rescue of the lifespan decrease by expressing transgenic *adr-1*, probably because of insufficient expression of the transgene in the germline and early embryonic cells because as the transgene is expressed from an extra chromosomal array that limits expression in the germline. This may also be the reason why transgenic ADR-1 with mutations in the dsRNA binding domains rescues to the same degree as the wild-type transgenic ADR-1. It is also possible that the different lifespan phenotypes in *adr-1* and *adr-2* are a result of distinct functions of these genes, which might not be related to RNA editing of one specific gene.

The protruding vulval (pvl) phenotype seems to be specific to *adr-1* mutants as suggested before (Tonkin et al., 2002), although in a very low penetrance. This phenotype was not allele specific as we observed it in two different alleles and we could partially rescue this phenotype with *adr-1* transgenes. Another phenotype that seems to be specific to *adr-1* mutants is the bag of worms (BOW) phenotypes when the mutant worms are subjected to *unc-22* RNAi. Both phenotypes, pvl and BOW, are probably related to each other. *adr-1*, *unc-22* and *unc-54* are expressed in the vulva (Moerman et al., 1988, Tonkin et al., 2002) and bind to each other. As *unc-22* and *unc-54* do not appear to be regulated by RNA editing, these phenotypes are probably not related to RNA editing and possibly ADR-1 has other functions not related to RNA editing, including regulating vulva formation.

In general *adr-1* mutant phenotypes are more severe than *adr-2* phenotypes and the double mutant phenotypes seems to be the middle ground between the phenotypes of the two single mutations for most phenotypes. Thus, it is likely that changes in the editing levels are more harmful than a complete loss of editing.

### RNA editing process is highly regulated

Our results show that ADR-1 has a significant effect on edited genes. We found that there is a decrease in the overall expression level of genes edited within 3’UTRs, that ADR-1 binds edited genes and that a significant portion of upregulated genes in *adr-1* mutants are genes edited at the L4 stage. Moreover, we found that the effect of ADR-1 on editing is stronger at the L4 developmental stage than at the embryo stage. Thus, ADR-1 can both upregulate and downregulate the expression of edited genes, and this might depend on the level of editing in a specific gene or even a specific site within a gene.

ADR-1 does not have deamination activity and was suggested to regulate editing by binding to ADR-2 targets and facilitating or preventing ADR-2 activity (Washburn et al., 2014). Indeed, we observed a very significant portion of ADR-1 binding targets are edited genes. The RIP-seq experiments that we performed do not exclude indirect binding to edited genes, however the RNA binding domains in the ADR-1 protein were shown to play an important role in editing regulation (Rajendren et al., 2018, Washburn et al., 2014). When we used a mutated version of ADR-1-FLAG with mutations in the RNA binding domain, the rescue of the vulva and lifespan phenotypes was similar to that of the strain expressing the wild-type ADR-1-FLAG transgene. Although Washburn et al (Washburn et al., 2014) demonstrated that ADR-1 with mutations in both dsRNA binding domains cannot bind several edited genes by RIP experiments coupled to qPCR, it is possible that the mutated ADR-1 lacks the ability to bind ADR-2 targets but can bind other mRNAs. Another hypothesis is that some of *adr-1* mutant phenotypes are not related to ADR-1 function in editing. Therefore, it is possible that the RNA binding domains are needed for ADR-1 function in editing and the inactive deamination domain has evolved to perform other ADR-1 functions such as in muscle formation.

By detecting edited sites that occur only in *adr-1* mutants, we found that the effect of ADR-1 on editing is more significant at the L4 developmental stage than in the embryo stage. This result goes well together with our findings that the expression of several edited genes is also upregulated in *adr-1* mutant worms (Figure 4) and that the phenotypes observed in *adr-1* mutants are associated with more advanced stages of development. These results suggest that the main function of ADR-1 is regulating editing by ADR-2 at L4 stage. We found that most of the editing sites that are unique to *adr-1* mutants at the embryo stage are in genes that undergo editing at other sites in wildtype worms. It is possible that these sites were not detected in wildtype worms because of the restriction of at least 5% editing in the analysis. It was shown that ADR-1 can both enhance and reduce the levels of editing (Washburn et al., 2014), therefore these sites might appear only in *adr-1* mutants because their editing level was enhanced enough to cross the threshold. This together with the absence of unique editing sites in the *adr-1* mutants leading to altered gene regulation, indicate that ADR-1 primarily promotes editing and does not prevent ADR-2 binding to specific sites and editing to result in altered gene expression.

Not many regulators of RNA editing are known in *C. elegans*. Only one protein other than ADR-1 was also shown to regulate editing. This protein, ADBP-1 (ADR-2 binding protein-1) was previously shown to alter ADR-2 nuclear localization (Ohta et al., 2008). We found that both ADR-2 and ADBP-1 regulate each other’s protein levels. It is not clear how ADR-2 affects ADBP-1, however it seems to be affected at the protein level, as the RNA expression of *adbp-1* is not affected in *adr-2* mutants. Other editing regulators might also be involved, regulating editing in a developmental-specific, tissue-specific and cellular-specific manner (Ganem and Lamm, 2017).

### The major role of RNA editing is to regulate RNA levels and not protein levels

In our previous study (Goldstein et al., 2017), we found that the level of expression in genes edited at their 3’UTR is slightly but significantly lower in worms mutated in *adr-1* and *adr-2* compared to wildtype. In this study, we also observed this reduction in gene expression in the single ADAR mutants at the embryo stage. Hundreds of genes were two-fold upregulated and downregulated at the RNA level in the single ADAR mutants and there was a significant portion of L4 edited genes in *adr-1* upregulated genes. However, only 22 genes had a significant change in protein levels in *adr-1* or *adr-2* mutant worms. From them only 4 genes were found to be edited (Goldstein et al., 2017), even though there was a high representation of edited genes in the proteomics data. Recently, RNA editing was shown to play an important part in suppressing the innate immune response in mammals (George et al., 2016, Liddicoat et al., 2015, Mannion et al., 2014) by marking self-produced dsRNA and by that preventing endogenous dsRNA from triggering the immune response. It is possible that RNA editing has a similar function in *C. elegans*, e.g. marking self dsRNA to prevent the immune response (RNAi) from processing and degrading the RNA (Reich et al., 2018, Ganem and Lamm, 2017). In addition, most of the edited sites in mammals and in *C. elegans* are in non-coding regions (for example, pseudogenes, intergenic regions and transposons) (Athanasiadis et al., 2004, Barak et al., 2009, Blow et al., 2004, Goldstein et al., 2017, Kim et al., 2004, Levanon et al., 2004, Li et al., 2009b, Warf et al., 2012, Wu et al., 2011). Thus, probably the main function of RNA editing in *C. elegans* is not to alter the content of the proteins in the cell but rather to buffer other processes such as RNAi.

## Experimental Procedures

### Maintenance and handling of *C. elegans* strains

The following strains were used in this study: Bristol N2, BB2 *adr-1 (gv6)* I, BB3 *adr-2 (gv42)* III, BB4 *adr-1 (gv6)* I; *adr-2 (gv42)* III) (Tonkin et al., 2002), BB19 *adr-1 (tm668)* I, RB886 *adr-2 (ok735)* III, BB21 *adr-1 (tm668)* I; *adr-2 (ok735)* III (Hundley et al., 2008); QD1 adbp-1(qj1) II (Ohta et al., 2008); HH76 a*dr-1*(tm668) I blmEx1[3XFLAG-*adr-1* genomic, *rab3*::*gfp*::*unc-54*]; HH116 *adr-1*(tm668) I blmEx1[3XFLAG-*adr-1* genomic with mutations in dsRBD1 (K223E, K224A, and K227A) and dsRBD2 (K583E, K584A and K587A), *rab3*::*gfp*::*unc-54*] HH134 a*dr-1*(tm668);*adr-2*(ok735) I blmEx1[3XFLAG-*adr-1* genomic, *rab3*::*gfp*::*unc-54*] (Washburn et al., 2014). All strains were grown at 20°C on NGM agar media with OP50 as food as described in (Brenner, 1974). The genotype of each of the strains was validated by PCR and sequencing before use (see supplementary materials). Embryos were isolated from adult worms by washing worms with M9 and sodium hypochloride. To collect L4 worms, embryos were left overnight in M9 in a nutator at 20°C and the hatched synchronized L1 larva were placed on NGM agar plate until they reached the L4 larva stage. L4 developmental stage was confirmed using binocular, by measuring ~650μm in length. Embryos or L4 larva worms were resuspended in either M9 for proteomic analysis or EN buffer for RNA extraction and frozen into pellets with liquid nitrogen. Only fluorescent worms were counted for every experiment that used strains HH76, HH116, and HH134.

### Life span assay

Synchronized L1 worms were plated and kept at 15°C, until they reached L4 stage (after about 48 hours). L4 worms from each tested strain were transferred to 5 FUdR plates, about 50 worms per plate. FUdR was added to the NGM agar before pouring the plates, to a final concentration of 4.95μM, in order to prevent the worms from having progeny. Live and dead worms were counted every 2-4 days. A worm was considered dead when it did not respond to touch of the platinum wire pick, and was subsequently removed from the plate. Worms that crawled over the edges of the plates and dried out were reduced from the total count. At least three biological replicates were performed for each experiment.

### Assays for Bag of worms Phenotype, vulva abnormalities, and hypersensitivity in worms

To perform RNAi, *unc-22* (Kamath et al., 2001) or lin-1 (Kamath et al., 2003) or empty L4440 plasmids were transformed into E. coli HT115 bacteria and were cultured overnight at 37°C in LB media containing 100 μg/ml ampicillin. The cultured bacteria were seeded onto *C. elegans* growth media plates (NGM) containing 100μg/ml ampicillin and 10 μg/ml IPTG and incubated overnight at RT overnight. Synchronized embryos were placed on the plates and incubated for 72-96 hours at 20°C before scoring. Worms were then scored using Nikon SMZ745 zoom stereomicroscope for PVL, BOW, or Multivulva phenotype. Worms presenting the phenotypes were counted in relative to the total number of worm in each plate. In the RNAi experiments the fraction of worms presenting the phenotype in the empty vector experiment was subtracted from the fraction of worms presenting the phenotype in the *unc-22* or *lin-1* RNAi experiments.

### mRNA-seq libraries preparation

Embryos and L4 worms frozen pellets were grounded to powder with a liquid nitrogen chilled mortar and pestle. RNA in high and low molecular weight fractions was extracted by mirVana kit (Ambion). RNA sequencing libraries were prepared from the high molecular weight fraction using the SMARTer^®^ Stranded RNA-Seq Kit (Clontech Laboratories) after ribosomal depletion by Ribozero kit (Epicenter) and sequenced with an Illumina HiSeq 2500. The sequence data from this study have been submitted to the NCBI Gene Expression Omnibus (GEO; http://www.ncbi.nlm. nih.gov/geo/ under accession number GSE110701.

### RNA editing sites and gene expression analysis

To extend the list of 3’UTR edited genes, non-repetitive edited sites found by (Goldstein et al., 2017) were reannotated according to the newest Wormbase (WS261). A gene was added to the list of 3’UTR edited genes if according to the new annotation it has multiple editing sites in its 3’UTR at the same orientation of the gene or multiple edited sites were found in proximity to the 3’UTR of the gene, less than 200bp apart, at the same orientation of the gene. At least three different biological RNA-seq samples were generated from N2 and ADAR mutant worms (BB2, BB19, BB3 and RB886), each at embryo and L4 stage. All reads were trimmed to 47nt and identical reads were merged. Sequences were aligned to gene transcripts from WS220 (Wormbase, www.wormbase.org) using Bowtie (Langmead et al., 2009). DESeq (Anders and Huber, 2010) package in R was used to identify differentially expressed genes. Significantly differently regulated genes were genes with at least 2-fold change in expression, with p-adj value ≤ 0.05. P-values for expression differences between 3’UTR edited genes and all genes were calculated by Welch two-sample t-test only for genes with p-adj value < 0.05. P-value for each mutant was calculated independently of other mutants. The Venn diagram overlap p-value was calculated by hypergeometric distribution using Phyper function in R. Identification of editing sites in *adr-1* mutants was done essentially as described in (Goldstein et al., 2017). The main difference is an increased stringency that a nucleotide change was selected only if it appeared in both *adr-1* mutants. In short, sequences from both *adr-1* mutants were aligned to WS220 genome using Bowtie (Langmead et al., 2009) with the restriction of not more than two alignments to exclude repetitive regions and were clustered using Samtools (Li et al., 2009a). Nucleotide change were selected if they appeared in both *adr-1* mutants, with at least 5% reads aligned to the site that contain the change and not more than 1% of reads with other nucleotide changes. Nucleotide changes were removed if they appeared in DNA-sequences or in RNA sequences from worms mutated in both ADAR genes, *adr-1* and *adr-2* (BB21 and BB4 strains).

### Proteomics

50μl of frozen pellets from three biological replicates of each strain in embryo or L4 stage were taken for the proteomics analysis. Proteins from the different samples were extracted by using urea buffer containing: 9M Urea, 400mM Ammonium bicarbonate [ABC] and 10mM DTT in the ratio of 600ul buffer to 50ul sample. The samples were then sonicated on ice (7’, 10s on/off pulse, 90% duty) and vortexed roughly. This procedure was repeated twice. The samples were then centrifuged at 14000rpm for 10’ and 17000g for 10’ in order to sediment the residual cuticle debris. Then the extracted proteins were trypsinized, and analyzed by LC-MS/MS on Q Exactive plus (Thermo). The data was analyzed with MaxQuant 1.5.2.8 vs the *Caenorhabditis elegans* part of the Uniprot database. Each mutant’s sample data was analyzed against the WT (N2) sample and known contaminants were removed. Only proteins that were identified with at least 2 peptides were tested for significant differences. Student t-test p-value threshold on LFQ intensities was set to 0.05. Difference threshold on LFQ intensities was set to +/-0.8.

### RIP-seq

Using worm strains containing a FLAG-ADR-1 transgene and lacking endogenous *adr-1* (HH76) or lacking both endogenous *adr-1* and *adr-2* (HH134) and *adr-1(-)* (BB21) worms as a negative control, the ADR-1 RNA immunoprecipitation was performed as previously described (Washburn et al., 2014). For two biological replicates of each RIP experiment, RNA extracted from portion of the input lysates and the immunoprecipitated RNA were subjected polyA selection using magnetic oligo-dt beads (Ambion) and a KAPA Stranded RNA-seq Library Preparation Kit (KAPA Biosystems). Equivalent amounts of the libraries were subjected to high-throughput sequencing on an Illumina NextSeq500 at the IU-Center for Genomics and Bioinformatics. The resulting 75 bp SE raw reads were trimmed of sequencing adapters, polyA tails, and repetitive elements using cutadapt (v1.9.1), and aligned with STAR (v2.4.0i) against RepBase (v18) to remove repetitive elements. Reads were then aligned to ce10 using the following STAR parameters: [outFilterMultimapNmax 10, outFilterScoreMinOverLread: 0.66, outFilterMatchN-minOverLread: 0.66, outFilterMismatchNmax: 10, outFilterMismatchNoverLmax: 0.3]. Read sorting and indexing was performed using Samtools 1.3.1. Gene expression was quantified with featureCounts (v1.5.0) using reads that map to exons. Raw read counts were input into DESeq2 (v1.18.1) to quantify differential expression for each IP/input pair using three replicates for the input samples and two replicates for the IP per condition. One IP replicate clustered independently by batch rather than genotype and therefore was removed from the final analysis. Genes enriched in IP were selected with a BH corrected p-value less than 0.05 and a log2 fold change greater than 0.5. Genes that are enriched in both ADR-1 samples (FLAG-ADR-1 in *adr-1(-)* and FLAG-ADR-1 in *adr-1(-);adr-2(-)*) and not in the negative control sample (*adr-1(-)*) were called as ADR-1 bound targets.

## Acknowledgements

We would like to thank the reviewers for their suggestions, which helped improve the manuscript. We thank Smoler proteomics center at the Technion for help with the proteomics analysis. Some strains were provided by the CGC, which is funded by NIH Office of Research Infrastructure Programs (P40 OD010440). This work was funded by The Israeli Centers of Research Excellence (I-CORE) program, (Center No. 1796/12 to ATL), The Israel Science Foundation (grants No. 644/13, 927/18 and 2154/16 to ATL), the Binational Israel-USA Science Foundation (grant No. 2015091 to ATL and HAH), the National Institutes of Health (F32GM119257-01A1 to SND) and the American Cancer Society (RSG-15-051 RMC to HAH).

## Author contributions

NSG, NBA, ACM, SND, MCW, OBZ, EM, HAH and ATL designed the experiments. NSG, NBA, ACM, SND, MCW, OBZ, and EM performed the experiments. NBA, ECW, GWY, and ATL performed the bioinformatics analysis. NSG, NBA, HAH, and ATL wrote the manuscript.

## References

Anders, S. & Huber, W. 2010. Differential expression analysis for sequence count data. Genome Biology, 11, R106.

Athanasiadis, A., Rich, A. & Maas, S. 2004. Widespread A-to-I RNA editing of Alu-containing mRNAs in the human transcriptome. PLoS Biol, 2, e391.

Barak, M., Levanon, E. Y., Eisenberg, E., Paz, N., Rechavi, G., Church, G. M. & Mehr, R. 2009. Evidence for large diversity in the human transcriptome created by Alu RNA editing. Nucleic Acids Res, 37, 6905–15.

Bass, B. L. 2006. How does RNA editing affect dsRNA-mediated gene silencing? Cold Spring Harb Symp Quant Biol, 71, 285–92.

Benian, G. M., Kiff, J. E., Neckelmann, N., Moerman, D. G. & Waterston, R. H. 1989. Sequence of an unusually large protein implicated in regulation of myosin activity in C. elegans. Nature, 342, 45–50.

Blow, M., Futreal, P. A., Wooster, R. & Stratton, M. R. 2004. A survey of RNA editing in human brain. Genome Res, 14, 2379–87.

Brenner, S. 1974. The genetics of Caenorhabditis elegans. Genetics, 77, 71–94.

Burns, C. M., Chu, H., Rueter, S. M., Hutchinson, L. K., Canton, H., Sanders-bush, E. & Emeson, R. B. 1997. Regulation of serotonin-2C receptor G-protein coupling by RNA editing. Nature, 387, 303–8.

Deffit, S. N., Yee, B. A., Manning, A. C., Rajendren, S., Vadlamani, P., Wheeler, E. C., Domissy, A., Washburn, M. C., Yeo, G. W. & Hundley, H. A. 2017. The C. elegans neural editome reveals an ADAR target mRNA required for proper chemotaxis. eLife, 6.

Ganem, N. S. & Lamm, A. T. 2017. A-to-I RNA editing - thinking beyond the single nucleotide. RNA biology, 0.

George, C. X., Ramaswami, G., Li, J. B. & Samuel, C. E. 2016. Editing of Cellular Self-RNAs by Adenosine Deaminase ADAR1 Suppresses Innate Immune Stress Responses. The Journal of Biological Chemistry, 291, 6158–6168.

Goldstein, B., Agranat-tamir, L., Light, D., Ben-naim Zgayer, O., Fishman, A. & Lamm, A. T. 2017. A-to-I RNA editing promotes developmental stage-specific gene and lncRNA expression. Genome Research, 27, 462–470.

Higuchi, M., Maas, S., Single, F. N., Hartner, J., Rozov, A., Burnashev, N., Feldmeyer, D., Sprengel, R. & Seeburg, P. H. 2000. Point mutation in an AMPA receptor gene rescues lethality in mice deficient in the RNA-editing enzyme ADAR2. Nature, 406, 78–81.

Hundley, H. A., Krauchuk, A. A. & Bass, B. L. 2008. C. elegans and H. sapiens mRNAs with edited 3′ UTRs are present on polysomes. RNA, 14, 2050–2060.

Kamath, R. S., Fraser, A. G., Dong, Y., Poulin, G., Durbin, R., Gotta, M., Kanapin, A., Bot, N. L., Moreno, S., Sohrmann, M., Welchman, D. P., Zipperlen, P. & Ahringer, J. 2003. Systematic functional analysis of the Caenorhabditis elegans genome using RNAi. Nature, 421, 231.

Kamath, R. S., Martinez-campos, M., Zipperlen, P., Fraser, A. G. & Ahringer, J. 2001. Effectiveness of specific RNA-mediated interference through ingested double-stranded RNA in Caenorhabditis elegans. Genome Biology, 2, research0002.1–research0002.10.

Kim, D. D., Kim, T. T., Walsh, T., Kobayashi, Y., Matise, T. C., Buyske, S. & Gabriel, A. 2004. Widespread RNA editing of embedded alu elements in the human transcriptome. Genome Res, 14, 1719–25.

Knight, S. W. & Bass, B. L. 2002. The role of RNA editing by ADARs in RNAi. Mol Cell, 10, 809–17.

Langmead, B., Trapnell, C., Pop, M. & Salzberg, S. L. 2009. Ultrafast and memory-efficient alignment of short DNA sequences to the human genome. Genome Biol, 10, R25.

Levanon, E. Y., Eisenberg, E., Yelin, R., Nemzer, S., Hallegger, M., Shemesh, R., Fligelman, Z. Y., Shoshan, A., Pollock, S. R., Sztybel, D., Olshansky, M., Rechavi, G. & Jantsch, M. F. 2004. Systematic identification of abundant A-to-I editing sites in the human transcriptome. Nat Biotechnol, 22, 1001–5.

Li, H., Handsaker, B., Wysoker, A., Fennell, T., Ruan, J., Homer, N., Marth, G., Abecasis, G., Durbin, R. & Genome project data processing, s. 2009a. The Sequence Alignment/Map format and SAMtools. Bioinformatics (Oxford, England), 25, 2078–2079.

Li, J. B., Levanon, E. Y., Yoon, J.-K., Aach, J., Xie, B., Leproust, E., Zhang, K., Gao, Y. & Church, G. M. 2009b. Genome-Wide Identification Of Human RNA Editing Sites by Parallel DNA Capturing and Sequencing. Science, 324, 1210–1213.

Liddicoat, B. J., Piskol, R., Chalk, A. M., Ramaswami, G., Higuchi, M., Hartner, J. C., Li, J. B., Seeburg, P. H. & Walkley, C. R. 2015. RNA editing by ADAR1 prevents MDA5 sensing of endogenous dsRNA as nonself. Science, 349, 1115–1120.

Mannion, N. M., Greenwood, S. M., Young, R., Cox, S., Brindle, J., Read, D., NellÅKer, C., Vesely, C., Ponting, C. P., Mclaughlin, P. J., Jantsch, M. F., Dorin, J., Adams, I. R., Scadden, A. D. J., Ohman, M., Keegan, L. P. & O’Connell, M. A. 2014. The RNA-editing enzyme ADAR1 controls innate immune responses to RNA. Cell Reports, 9, 1482–1494.

Moerman, D. G., Benian, G. M., Barstead, R. J., Schriefer, L. A. & Waterston, R. H. 1988. Identification and intracellular localization of the unc-22 gene product of Caenorhabditis elegans. Genes & Development, 2, 93–105.

Moerman, D. G., Plurad, S., Waterston, R. H. & Baillie, D. L. 1982. Mutations in the unc-54 myosin heavy chain gene of Caenorhabditis elegans that alter contractility but not muscle structure. Cell, 29, 773–781.

Morse, D. P., Aruscavage, P. J. & Bass, B. L. 2002. RNA hairpins in noncoding regions of human brain and Caenorhabditis elegans mRNA are edited by adenosine deaminases that act on RNA. Proc Natl Acad Sci U S A, 99, 7906–11.

Morse, D. P. & Bass, B. L. 1999. Long RNA hairpins that contain inosine are present in Caenorhabditis elegans poly(A)+ RNA. Proc Natl Acad Sci U S A, 96, 6048–53.

Ohta, H., Fujiwara, M., Ohshima, Y. & Ishihara, T. 2008. ADBP-1 regulates an ADAR RNA-editing enzyme to antagonize RNA-interference-mediated gene silencing in Caenorhabditis elegans. Genetics, 180, 785–796.

Palladino, M. J., Keegan, L. P., O’Connell, M. A. & Reenan, R. A. 2000. A-to-I pre-mRNA editing in Drosophila is primarily involved in adult nervous system function and integrity. Cell, 102, 437–449.

Pullirsch, D. & Jantsch, M. F. 2010. Proteome diversification by adenosine to inosine RNA editing. RNA biology, 7, 205–212.

Rajendren, S., Manning, A. C., Al-awadi, H., Yamada, K., Takagi, Y. & Hundley, H. A. 2018. A protein-protein interaction underlies the molecular basis for substrate recognition by an adenosine-to-inosine RNA-editing enzyme. Nucleic Acids Research.

Reich, D. P., Tyc, K. M. & Bass, B. L. 2018. C. elegans ADARs antagonize silencing of cellular dsRNAs by the antiviral RNAi pathway. Genes & Development, 32, 271–282.

Sakurai, M., Shiromoto, Y., Ota, H., Song, C., Kossenkov, A. V., Wickramasinghe, J., Showe, L. C., Skordalakes, E., Tang, H.-Y., Speicher, D. W. & Nishikura, K. 2017. ADAR1 controls apoptosis of stressed cells by inhibiting Staufen1-mediated mRNA decay. Nature Structural & Molecular Biology, advance online publication.

Sebastiani, P., Montano, M., Puca, A., Solovieff, N., Kojima, T., Wang, M. C., Melista, E., Meltzer, M., Fischer, S. E. J., Andersen, S., Hartley, S. H., Sedgewick, A., Arai, Y., Bergman, A., Barzilai, N., Terry, D. F., Riva, A., Anselmi, C. V., Malovini, A., Kitamoto, A., Sawabe, M., Arai, T., Gondo, Y., Steinberg, M. H., Hirose, N., Atzmon, G., Ruvkun, G., Baldwin, C. T. & Perls, T. T. 2009. RNA editing genes associated with extreme old age in humans and with lifespan in C. elegans. PloS One, 4, e8210.

Simmer, F., Tijsterman, M., Parrish, S., Koushika, S. P., Nonet, M. L., Fire, A., Ahringer, J. & Plasterk, R. H. A. 2002. Loss of the putative RNA-directed RNA polymerase RRF-3 makes C. elegans hypersensitive to RNAi. Current biology: CB, 12, 1317–1319.

Solomon, O., Oren, S., Safran, M., Deshet-unger, N., Akiva, P., Jacob-Hirsch, J., Cesarkas, K., Kabesa, R., Amariglio, N., Unger, R., Rechavi, G. & Eyal, E. 2013. Global regulation of alternative splicing by adenosine deaminase acting on RNA (ADAR). RNA (New York, N.Y.), 19, 591–604.

Tonkin, L. A. & Bass, B. L. 2003. Mutations in RNAi rescue aberrant chemotaxis of ADAR mutants. Science 302, 1725.

Tonkin, L. A., Saccomanno, L., Morse, D. P., Brodigan, T., Krause, M. & Bass, B. L. 2002. RNA editing by ADARs is important for normal behavior in Caenorhabditis elegans. EMBO J, 21, 6025–35.

Wang, Q., Khillan, J., Gadue, P. & Nishikura, K. 2000. Requirement of the RNA editing deaminase ADAR1 gene for embryonic erythropoiesis. Science (New York, N.Y.), 290, 1765–1768.

Warf, M. B., Shepherd, B. A., Johnson, W. E. & Bass, B. L. 2012. Effects of ADARs on small RNA processing pathways in C. elegans. Genome Res, 22, 1488–98.

Washburn, M. C., Kakaradov, B., Sundararaman, B., Wheeler, E., Hoon, S., Yeo, G. W. & Hundley, H. A. 2014. The dsRBP and inactive editor, ADR-1, utilizes dsRNA binding to regulate A-to-I RNA editing across the C. elegans transcriptome. Cell reports, 6, 599–607.

Waterston, R. H. 1989. The minor myosin heavy chain, mhcA, of Caenorhabditis elegans is necessary for the initiation of thick filament assembly. The EMBO journal, 8, 3429–3436.

Werry, T. D., Loiacono, R., Sexton, P. M. & Christopoulos, A. 2008. RNA editing of the serotonin 5HT2C receptor and its effects on cell signalling, pharmacology and brain function. Pharmacol Ther, 119, 7–23.

Whipple, J. M., Youssef, O. A., Aruscavage, P. J., Nix, D. A., Hong, C., Johnson, W. E. & Bass, B. L. 2015. Genome-wide profiling of the C. elegans dsRNAome. RNA (New York, N.Y.), 21, 786–800.

Wu, D., Lamm, A. T. & Fire, A. Z. 2011. Competition between ADAR and RNAi pathways for an extensive class of RNA targets. Nat Struct Mol Biol, 18, 1094–1101.

Zhao, H.-Q., Zhang, P., Gao, H., He, X., Dou, Y., Huang, A. Y., Liu, X.-Q., Ye, A. Y., Dong, M.-Q. & Wei, L. 2015. Profiling the RNA editomes of wild-type C. elegans and ADAR mutants. Genome Research, 25, 66–75.

